# Temporally organized activity in mouse V1 encodes newly sampled visual content during free movement

**DOI:** 10.64898/2026.08.21.746335

**Authors:** Jhoseph Shin, Elliott T. T. Abe, Philip R. L. Parker, Dylan M. Martins, Cristopher M. Niell

**Author notes:** Lead Contact / Correspondence: Cristopher M. Niell.

## Abstract

Natural vision is continuously shaped by an animal’s own movements, which determine what enters the visual system and when visual input changes. Gaze shifts are known to initiate a temporally structured sequence of activity in primary visual cortex (V1), but how the visual content sampled by each movement contributes to this sequence has remained unclear. We recorded visual input, eye and head movements, and V1 activity in freely moving mice, and asked how the visual content sampled on each gaze shift shapes the response. The magnitude of visual change induced by each gaze shift scaled the amplitude of responses according to each neuron’s characteristic response profile, while movement amplitude alone did not reproduce this modulation in darkness, supporting a role of visual input in driving the sequence. Activity following gaze shifts reflected each neuron’s spatial receptive field structure, and visual filters estimated under head-fixed conditions predicted the relative timing of spike responses during gaze shifts of freely moving animals, demonstrating that gaze shift responses encode visual information. At the population level, decoded V1 activity shifted toward the scene sampled after each gaze shift. Thus, across single-neuron, spatial, temporal, and population measures, V1 activity tracked the content of each sample beyond movement timing alone, indicating that gaze shifts act as sampling events that result in V1 encoding visual information in temporally ordered responses.

**HIGHLIGHTS:**

- Gaze shifts result in rapid shifts in the visual input and evoke a temporal sequence of responses in V1
- The magnitude of visual change across each gaze shift scales the strength of V1 responses
- Spatial receptive field structure is preserved in the activity following gaze shifts
- Visual tuning measured during head-fixed conditions predicts the timing of gaze shift responses during free movement
- Together, these results show that gaze shifts temporally organize V1 coding of newly sampled visual input

## INTRODUCTION

During natural behavior, the visual system processes input that is actively sampled by the animal’s own movements (Yarbus 1967; Land and Hayhoe 2001; Hayhoe and Ballard 2005). Each time an animal shifts its gaze, a new region of the world is swept across the retina, delivering a burst of visual input that the brain must rapidly process and interpret (Kuang et al. 2012; Rucci and Victor 2015). The timing and consequences of these movements structure the visual input available to the brain over time (Ahissar and Arieli 2001; Schroeder et al. 2010). In primates, saccadic eye movements are tightly coordinated with neural activity across visual and memory areas; neurons in visual cortex show modulated responses around saccade or fixation onsets (Leopold and Logothetis 1998; Rajkai et al. 2008), while hippocampal oscillations are reset by eye movements during visual exploration (Jutras et al. 2013). More broadly, the rhythmic cycle of redirecting and stabilizing gaze plays an active role in how visual information is sampled, segmented, and encoded over time (Boi et al. 2017; Ahissar and Arieli 2001; Schroeder et al. 2010; Samonds et al. 2018).

Although rodents lack a fovea, freely moving mice also perform coordinated head and eye movements (Meyer et al. 2020; Michaiel et al. 2020). These movements fall into two distinct modes: gaze shifting and compensatory movements. During gaze shifts, the head and eyes move together in the same direction, rapidly redirecting where the animal is looking and delivering new visual input to the retina (Meyer et al. 2020; Michaiel et al. 2020). During compensatory eye movements, the eyes move in the direction opposite to the head, stabilizing the current view as the head rotates (Meyer et al. 2020; Michaiel et al. 2020). This alternation between redirecting and stabilizing gaze is a pattern seen across many species known as “saccade-and-fixate” behavior (Land 2019). In freely moving mice, this behavioral structure shapes where visual input is sampled and how it changes over time on the retina, while V1 activity also reflects eye and head position (Meyer et al. 2020; Michaiel et al. 2020; Parker et al. 2022).

Recent work using a system that simultaneously records neural activity, eye position, head movement, and the visual scene in freely moving mice has begun to characterize how V1 responds during natural behavior (Parker et al. 2022). This approach allows visual receptive fields to be mapped in a freely moving animal, and has shown that many V1 neurons are influenced by both the visual scene and the animal’s eye and head position during free movement (Parker et al. 2022). Using this recording technique, we recently showed that each gaze shift triggers a structured sequence of V1 activity that unfolds over hundreds of milliseconds (Parker et al. 2023). When neurons were grouped by their gaze shift response profiles, the groups included cells that responded with an increase in activity of either short or long latency, cells that first decreased and then increased their firing, and cells that showed sustained decreases in firing. Following each gaze shift, these groups activate in a stereotyped temporal order, forming a temporal progression across the population that we refer to as the gaze shift response sequence. This sequence was largely absent when animals moved in darkness, indicating that visual input is necessary for its expression. Related gaze behavior has been characterized in freely moving marmosets using head-mounted eye tracking (Singh et al. 2025), and similar gaze-related V1 dynamics have been observed in both head-fixed freely gazing marmosets (Parker et al. 2023) and freely moving marmosets (Li et al. 2026), suggesting that this organization of visual responses around gaze shifts may be a shared feature across the mammalian visual cortex.

These findings showed that visual input is necessary for the expression of the gaze shift response sequence and that response timing varies with spatial frequency preference. However, how the specific visual input sampled on individual gaze shifts contributes to this sequence, and whether it accounts for event-to-event variation in V1 activity, has remained unclear. This question is important because V1 activity during natural behavior reflects widespread movement-related signals (Stringer et al. 2019; Musall et al. 2019), yet a growing body of work suggests that much of this activity may reflect the visual input those movements generate rather than the movements themselves (Talluri et al. 2023). Individual gaze shifts offer a way to dissociate the effects of movement from those of the visual input it generates within a single behavior, since each event delivers both a movement and a new visual sample.

Here, we combined measurements of visual input, eye position, head movement, and V1 activity in freely moving mice to test how the content sampled by individual gaze shifts shapes cortical responses. Across four independent analyses examining response magnitude, spatial receptive field structure, response timing, and population decoding, our results supported the conclusion that V1 activity around each gaze shift is strongly organized by the visual content that movement brings into view.

## RESULTS

### Gaze shifts produce rapid visual changes and structured V1 responses

We first characterized how gaze shifts change visual input during free movement. Mice explored an open arena while we recorded head position, eye position, the visual scene from a head-mounted “world camera,” and V1 spiking activity (Figure 1A). Because the world camera was fixed to the head rather than the eye, the raw camera movie does not directly correspond to the visual input falling on the retina. We therefore used the shift correction procedure of Parker et al. (2022) to estimate retinal input from the world camera movie. This correction incorporates measured head and eye positions and was used for all subsequent visual input analyses.

**Figure 1.**
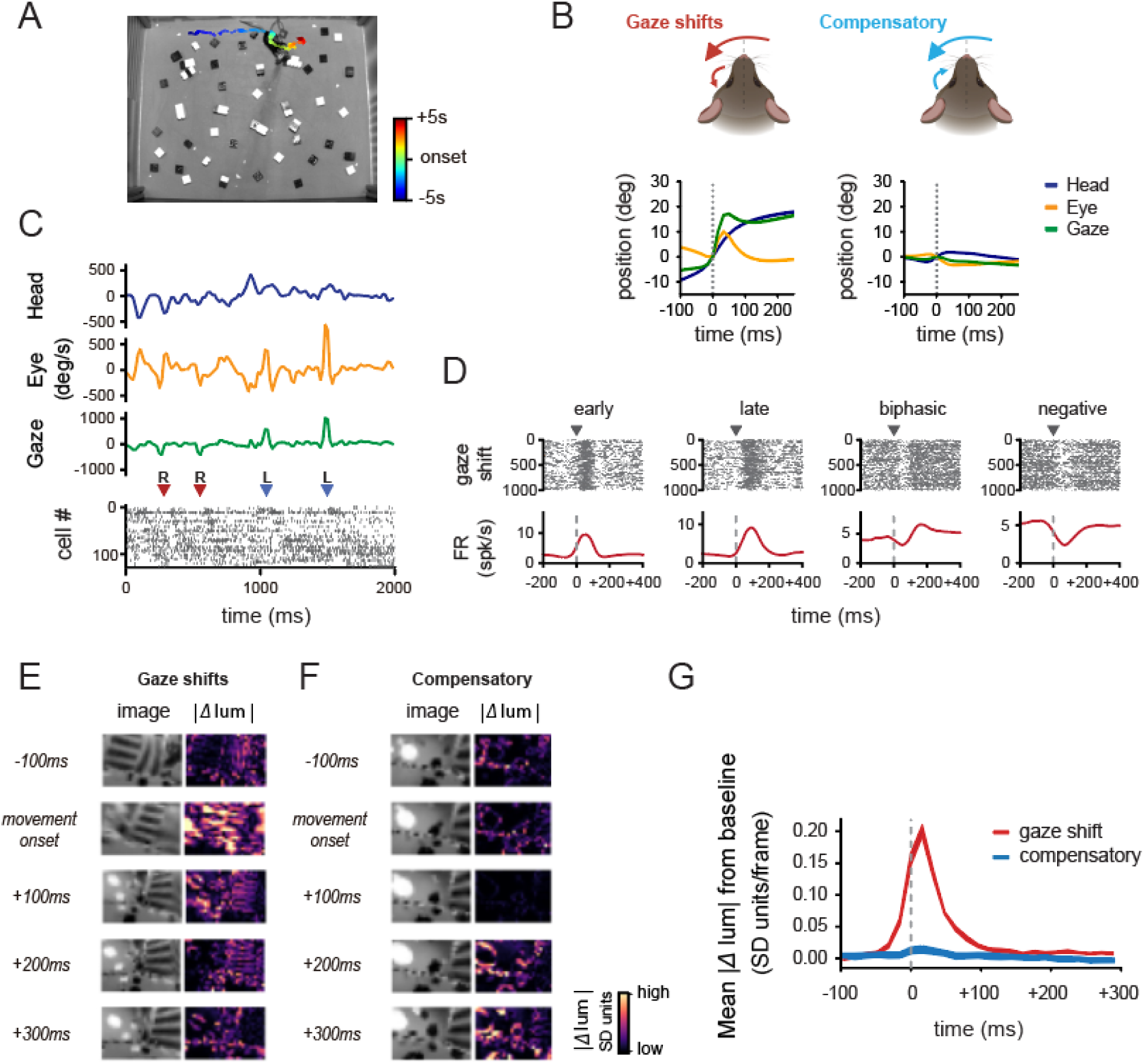
Gaze shifts rapidly change visual input during free movement. (**A**) Top down view of the experimental arena with the animal’s trajectory aligned to an example gaze shift onset. (**B**) Schematic and average head, eye, and gaze trajectories for gaze shifts and compensatory movements. Gaze was defined as head plus eye position. Dashed lines indicate movement onset; traces show the mean across animals (n = 8). (**C**) Example head, eye, and gaze angular velocity traces with simultaneously recorded V1 spiking activity. Units are sorted by positive response latency relative to gaze shift onset. Triangles mark detected gaze shifts and their direction (red = right, blue = left). (**D**) Gaze shift aligned spike rasters and firing rate traces for early, late, biphasic, and negative V1 response clusters. (**E**) Example image frames and luminance change maps (|*Δ*lum|) aligned to gaze shift onset. Δlum indicates the magnitude of pixel-wise changes in standardized luminance between consecutive frames. (**F**) Same as (E), aligned to compensatory movement onset. (**G**) Mean |*Δ* lum| from baseline aligned to gaze shift and compensatory movement onset. Values were baseline-subtracted using the −500 to −100 ms period. Lines and shading indicate mean ± SEM across animals (n = 8).

As previously described, we classified movements into two main categories based on the relative direction of head and eye movements: gaze shifts or compensatory movements (Figure 1B). During gaze shifts, the head and eyes moved in the same direction, producing a rapid change in gaze position relative to the external world. During compensatory movements, the eyes moved opposite to the head, reducing the net displacement of gaze. Consistent with this definition, gaze shifts produced larger changes in gaze direction than compensatory movements (median: 18.23° vs. 1.85°; Wilcoxon rank-sum test, p < 0.0001). During locomotion, mice generated 104.2 ± 6.13 gaze shifts per minute on average. Head and eye movements during gaze shifts also showed stereotyped temporal profiles, with peak head and eye velocities of 198.79 ± 5.18°/s and 368.82 ± 19.67°/s, respectively (mean ± SEM; Figure 1B and Figure S1).

Coordinated head and eye movements produced rapid shifts in gaze position, and V1 units showed event-aligned responses with different response latencies (an example is shown in Figure 1C). To characterize the diversity of V1 responses to gaze shifts, we grouped gaze-responsive neurons based on their event-aligned firing profiles using PCA followed by k-means clustering, identifying four response clusters: early cells (n = 51), late cells (n = 77), biphasic cells (n = 90), and negative cells (n = 52; total n = 270 of 649 cells from 8 mice; Figure 1D). Units were included if they met the firing rate and gaze shift modulation criteria described in Methods. Unless otherwise noted, we used these same cells throughout the study. These clusters captured the main temporal profiles observed around gaze shift onset.

The nature of the visual change depended on movement type. In an example image sequence, gaze shifts rapidly altered the estimated retinal input (Figure 1E). To quantify how gaze shifts changed the estimated retinal input, we computed luminance change maps (*Δ* lum) as the magnitude of frame-to-frame changes in pixel-wise z-scored luminance from the shift-corrected world camera video. Around gaze shift onset, these maps showed large local luminance changes spread across the image. By contrast, compensatory movements showed weaker luminance changes around movement onset (Figure 1F), consistent with the smaller net gaze displacement during these events. This difference was also evident across animals: gaze shifts produced a rapid transient increase in mean absolute image change around movement onset that was greater than that observed during compensatory movements over the 0-100 ms period (paired Wilcoxon signed-rank test, p = 0.0078, n = 8 mice; Figure 1G).

These observations established two key features of gaze shifts during free movement: they rapidly altered the visual input available to the animal, and they were accompanied by a set of distinct temporal response profiles in V1. We then asked whether differences in the visual content sampled on individual gaze shifts explain how these responses vary across events.

### Visual change shapes temporal profile of gaze shift responses

We quantified visual change across individual gaze shifts and asked whether the amount of change shaped how strongly V1 neurons responded and whether their firing increased or decreased. Because natural environments are not uniform, different gaze shifts will bring different amounts of new visual content into view, and we hypothesized that if the visual content sampled on each event shapes gaze shift responses, responses should also vary with the degree of visual change across events.

We computed a luminance change index as the mean absolute value of the pixel-wise change in luminance across the image, between consecutive frames of the shift-corrected world camera video. We also computed a spatial frequency change index as the change in spectral power in the image within three spatial frequency ranges. To determine the change for each gaze shift, we took the peak of each measure after gaze shift onset, standardized them across events within each session, and averaged the two values to obtain one index per event. Events were divided into high and low visual change groups using a median split within each session. Because the index is defined from these two features, high visual change events showed larger luminance and spatial frequency transients around gaze shift onset by construction (Figure 2A and 2B). Interestingly, these differences in visual input were reflected in V1 activity: high visual change gaze shifts resulted in larger amplitude responses, both positive and negative (Figure 2C-E). Representative neurons showed corresponding differences in their responses across the two conditions (Figure 2C), and their differences were also present in the population activity (Figure 2D). This shows that V1 responses varied not only with the occurrence of a gaze shift, but also with the magnitude of the visual transients accompanying that movement.

**Figure 2.**
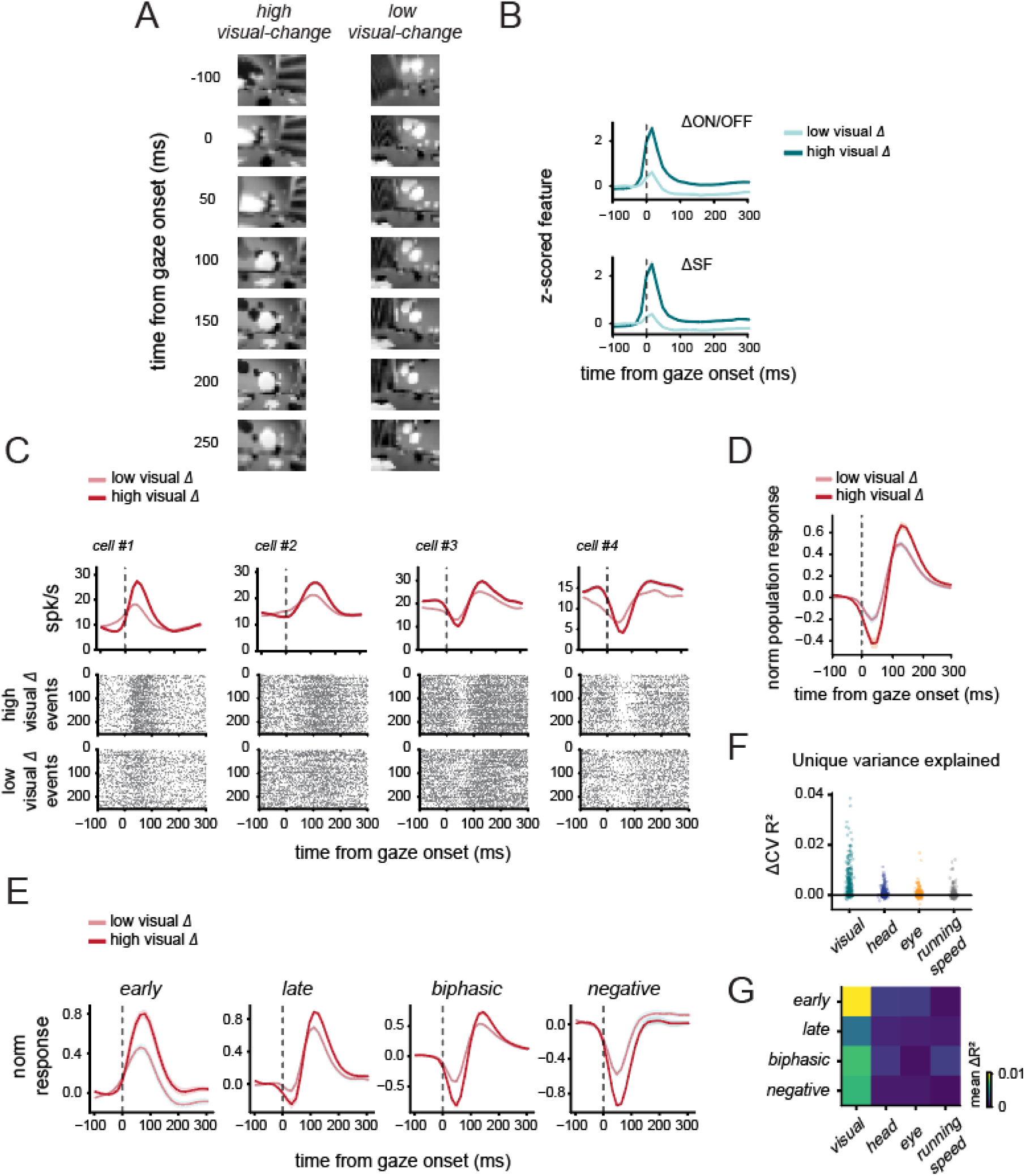
Visual transients during gaze shifts shape V1 population responses. (**A**) Example image sequences for gaze shifts with high or low visual change. (**B**) Mean ΔON/OFF and *Δ*SF feature timecourses for low and high visual change events. (**C**) Example neurons showing gaze shift responses that scale with visual change. Traces show mean firing rates; rasters show event-aligned spikes. (**D**) Mean normalized V1 population responses for low and high visual change gaze shifts. (**E**) Cluster-averaged responses for early (n = 51), late (n = 77), biphasic (n = 90), and negative (n = 52) cells. (**F**) Per cell unique variance explained by visual, head, eye, and running speed predictors, measured as *Δ*CV R² across V1 neurons (n = 270 cells, 8 mice). (**G**) Cluster-averaged unique ΔCV R² for each predictor.

To determine how this population-level difference was expressed across the different response clusters, we next examined high and low visual change responses separately for each cluster. The effect differed across clusters in a way that reflected each cluster’s characteristic response profile. Early and late cells showed larger positive responses during high visual change events, whereas biphasic and negative cells showed deeper negative responses (Figure 2E). This pattern was not consistent with a uniform population-wide increase in firing. Instead, visual change shaped responses in different directions across clusters, following the response profile already expressed by each group.

We next used the visual transient index as a continuous variable across gaze shift events, and asked whether it explained response variability independently of the accompanying motor signals, since gaze shifts with larger visual changes were also associated with larger head, eye, and gaze amplitudes (Figure S2A). For each neuron, we estimated the unique variance explained by the visual transient index, head amplitude, eye amplitude, and running speed using cross-validated regression (Figures 2F and 2G). The visual transient index explained more unique variance than any motor variable, with the largest contributions to the positive responses of early and late cells and the negative responses of biphasic and negative cells (all q < 3.5 × 10⁻⁴; Figure 2G). Visual transient magnitude was also the most frequent dominant predictor in every cluster (55% to 71% of cells; Figures 2G, S2B and S2C). The absolute explained variance was modest, as expected due to the brief event windows that typically contained few spikes.

To examine how visual change shaped each neuron’s response over time, we measured how firing rate changes covaried with the visual transient index across events at each time bin around gaze shift onset. Figure 3A shows two example neurons at a given time point: greater visual change was associated with increased firing in the positive-responding neuron and decreased firing in the negative-responding neuron. We used the slope of this relationship as a measure of the impact of visual change on firing on a moment-to-moment basis. The resulting slopes followed each cluster’s temporal response profile: positive slopes aligned with firing rate increases in early and late cells, whereas negative slopes aligned with the ‘suppressive’ components of biphasic and negative cells (Figure 3B and 3C; Kruskal-Wallis positive: p = 6.4 × 10⁻²²; negative: p = 2.5 × 10⁻²¹). This indicates that larger visual changes amplified each cluster’s temporal profile rather than producing a uniform additive shift in firing.

**Figure 3.**
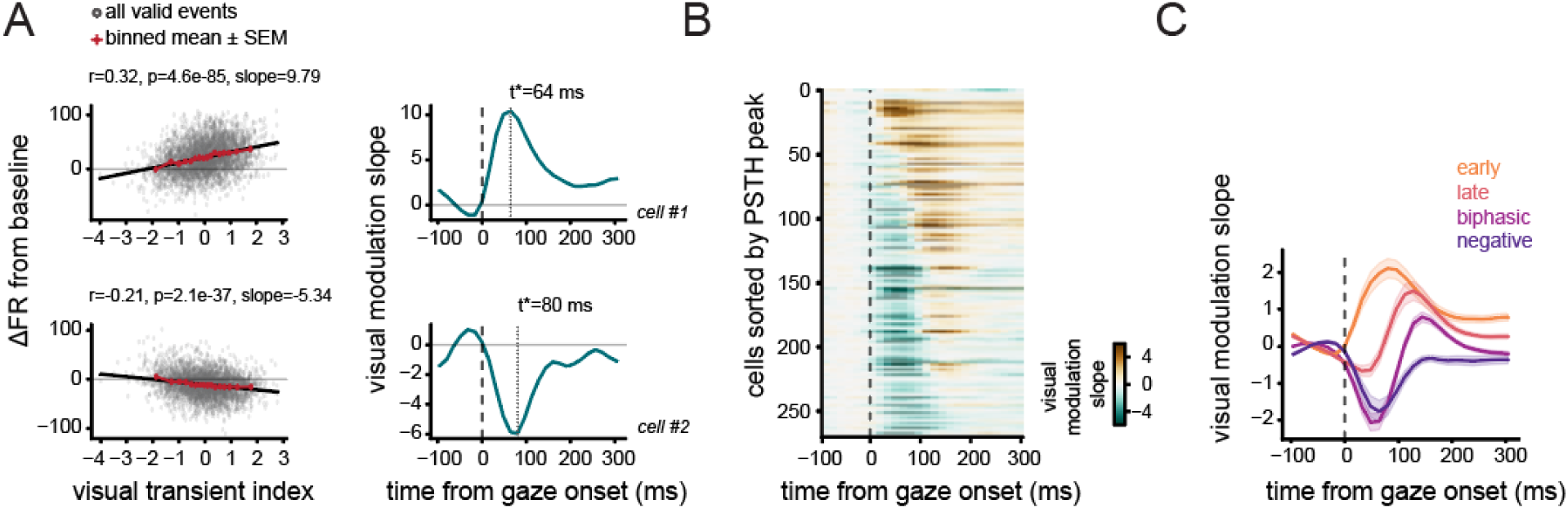
Event-by-event visual transients shape structured V1 response dynamics. (**A**) Example V1 neurons showing event-by-event relationships between post-gaze visual transient magnitude and baseline-subtracted firing rate changes. Right, visual modulation slope for the same neurons. The dashed line indicates the gaze shift onset. (**B**) Visual modulation slopes across V1 neurons (n = 270 cells, 8 mice), sorted by positive PSTH peak latency. Positive (*brown*) and negative values (*green*) indicate visual-transient-associated increases or decreases in ΔFR, respectively. (**C**) Cluster-averaged slopes for early (n = 51), late (n = 77), biphasic (n = 90), and negative (n = 52) response clusters. Visual transients preferentially enhanced positive responses in early/late cells and negative responses in biphasic/negative cells.

These analyses suggested that visual change was closely linked to evoking each cluster’s temporal profile, even after accounting for measured motor variables (Figure 2F and 2G). However, during natural gaze shifts, larger movements also tend to produce larger changes in the image, making it difficult to separate visual and motor contributions using light session data alone. We therefore analyzed a separate group of mice with paired light and dark recordings (n = 8), which allowed us to remove visual input while preserving gaze shift movements. We then asked whether movement amplitude alone could reproduce the cluster-specific modulation observed in light sessions. Cluster identity and preferred gaze direction were defined from the light session and applied to the corresponding dark session. In the dark session, splitting gaze shift events by total movement amplitude separated the movement traces, but produced little separation in cluster-averaged V1 responses (Figure S3A, B). Across sessions, the mean response difference between high- and low-amplitude events over 0 to 250 ms after gaze shift onset did not differ significantly from zero for any cluster type (early: ΔR = 0.004 ± 0.039, p = 0.775; late: 0.025 ± 0.072, p = 0.479; biphasic: −0.001 ± 0.054, p = 0.968; negative: −0.038 ± 0.085, p = 0.735; two-sided sign-flip tests). We also found a similar result using z-scored gaze amplitude rather than median split for each cluster (mean β over 0 to 250 ms: early = 0.001 ± 0.137, p = 1.000; late = −0.073 ± 0.080, p = 0.421; biphasic = −0.176 ± 0.124, p = 0.227; negative = −0.003 ± 0.066, p = 0.984; two-sided sign-flip tests; Figure S3C). These results indicate that visual change is the major factor that is linked to evoking each cluster’s temporal profile, rather than movement alone.

### Gaze shift responses reflect spatial receptive field structure

The previous analyses established that the visual input from each gaze shift shapes each neuron’s temporal profile. However, these analyses did not determine whether the elicited spikes preserved each neuron’s spatial visual selectivity, so we next asked whether the receptive structures are maintained during gaze shifts.

To estimate receptive fields from spikes during gaze shifts, we used a regularized linear model, fit separately to different recording periods: head-fixed white noise stimulus presentation, active periods of free movement, gaze shift epochs, and compensatory movement epochs. Gaze shift and compensatory receptive fields were estimated from equal numbers of timepoints. Then, we asked whether each estimated spatial pattern was reproducible across independent data splits. Receptive fields were estimated for training and test data separately, and a receptive field was classified as significant if its structure exceeded that of temporally shifted data (see Methods).

As expected, most of these cells showed significant receptive field structure during head-fixed white noise stimuli (245 of 270 cells, 90.7%; Figure 4A and 4C). Significant receptive field structures were also detected in most cells during active freely moving behavior (197 of 270 cells, 73.0%) and during gaze shift epochs (228 of 270 cells, 84.4%; Figure 4B and 4C). Compensatory movement epochs also contained spatially organized responses in a subset of cells, but this structure was detected less often than during gaze shifts (139 of 270 cells, 51.5%; Figure 4C). This difference was consistent across sessions: gaze shift epochs showed a higher fraction of cells with significant receptive field structure than frame-matched compensatory epochs by a median of 32.9 percentage points across sessions (n = 8 sessions; two-sided paired Wilcoxon signed-rank test, p = 0.0078). The same pattern was observed when the analysis was restricted to cells with significant head-fixed white noise receptive fields, where 210 of 245 cells (85.7%) showed significant receptive field structure during gaze shift epochs, compared with 126 of 245 cells (51.4%) during compensatory epochs.

**Figure 4.**
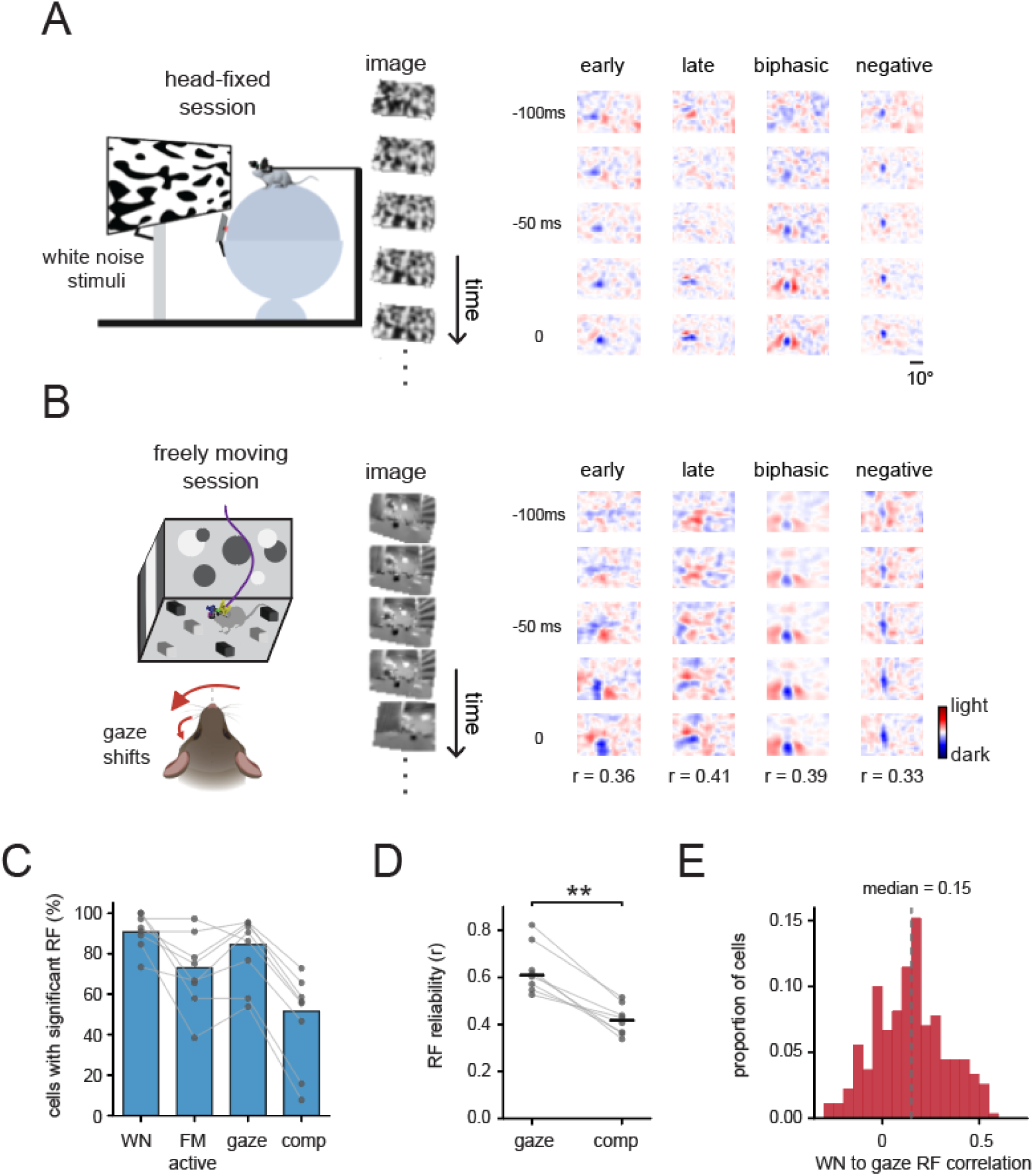
Gaze shift responses reflect spatial receptive field structure. (**A**) Head-fixed white noise stimulus schematic and example receptive field estimates from representative response classes. (**B**) Freely moving gaze shift schematic and gaze shift receptive field estimates from the same response classes. Values indicate spatial correlations between head-fixed white noise and gaze shift receptive fields. (**C**) Percentage of cells with significant receptive field structure in head-fixed white noise, active freely moving, gaze shift, and frame matched compensatory conditions. Significance was assessed using split-half receptive field reliability with temporally shifted control maps. Gray points and lines show session level values. (**D**) Session level split-half receptive field reliability during gaze shift and compensatory movement epochs. Each point shows the median training to test receptive field correlation across cells from one session, with paired values connected by lines. (**E**) Distribution of spatial correlations between head-fixed white noise and gaze shift receptive fields. The dashed line indicates the median.

As a complementary measure of receptive field reliability, we also computed correlations between the receptive fields estimated from the test and training dataset from gaze shift and compensatory movement epochs. This analysis treated reliability as a continuous measure rather than classifying each cell as significant or non-significant. For each session, test-train receptive field reliability was summarized as the median value of training vs test receptive field correlation across cells. Gaze shift epochs showed higher reliability than compensatory epochs across sessions (0.644 for gaze shifts and 0.419 for compensatory movements; p = 0.0078; Figure 4D). Thus, although compensatory epochs also contained measurable spatial structure, gaze shifts displayed spatially organized visual selectivity more robustly.

Lastly, we asked whether receptive fields estimated during gaze shifts were spatially similar to receptive fields measured independently during head-fixed white noise stimulation. Across the 270 cells, spatial correlation between head-fixed white noise and gaze shift receptive fields showed a positive median (median r = 0.154; Figure 4E). Thus, responses after gaze shifts reflected a reliable receptive field structure and matched each neuron’s receptive field estimated independently from the head-fixed condition. Together, these results indicate that gaze shift responses do not simply reflect a global modulation of V1 activity, but encode information about spatial structure in the visual input.

### Visual filters estimated under head-fixed conditions predict gaze shift response timing

The previous results show that gaze shift responses reflected spatial receptive field structure, but it remained unclear whether visual response properties measured under head-fixed conditions could account for their temporal structure during free movement. To test this, we trained feature-based visual filters from head-fixed sparse noise responses and applied them directly to shift-corrected world camera images collected during the freely moving session (Figure 5A; Figure S5). Because the model was not fit to gaze shift activity, any correspondence between predicted and actual responses would indicate that visual tuning measured under controlled conditions generalizes to the temporal organization of responses during natural behavior.

**Figure 5.**
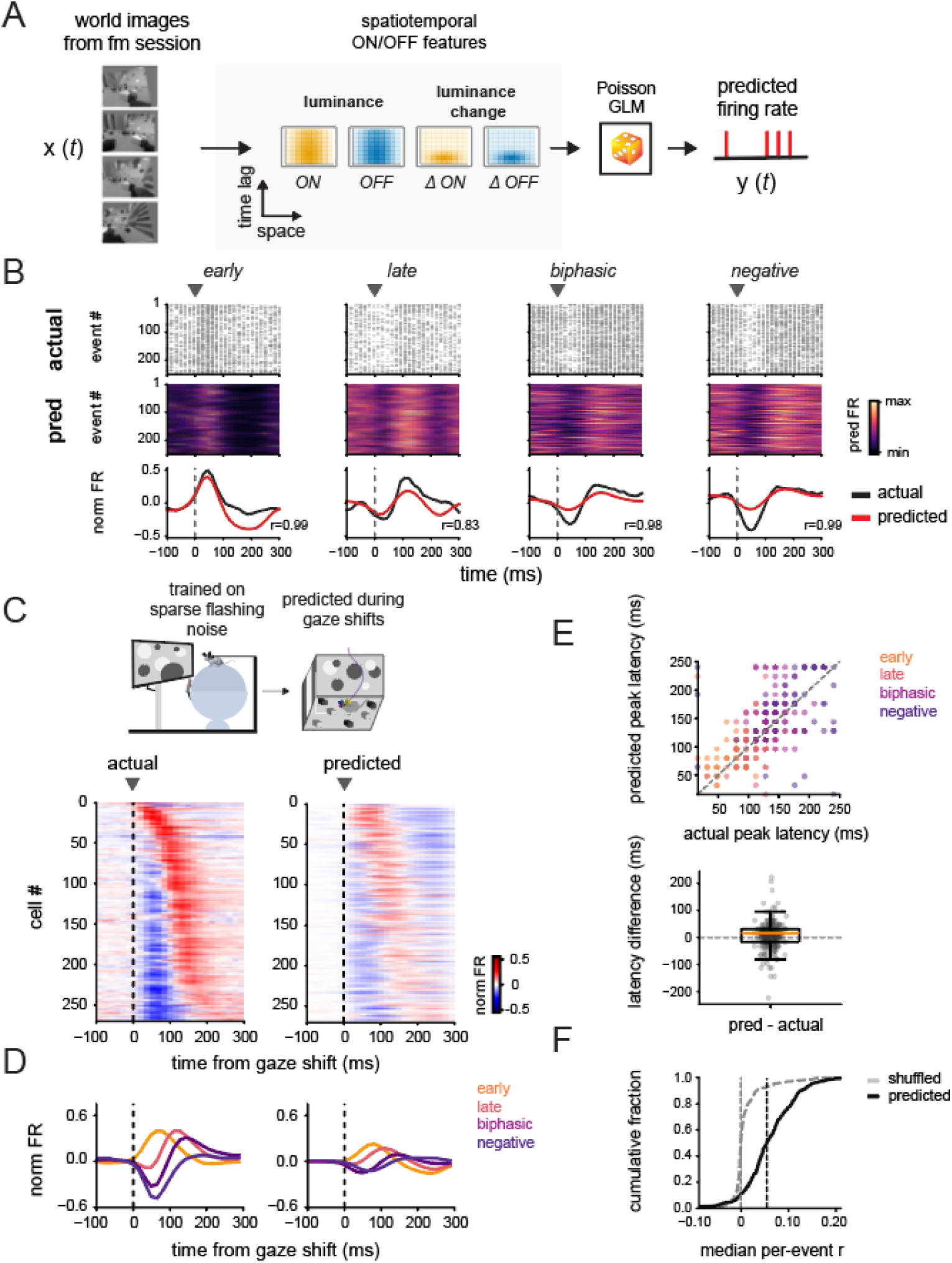
Head-fixed visual filters predict the timing of gaze shift responses. (A) Visual filters estimated from head-fixed sparse noise responses were applied directly to shift-corrected world camera frames during freely moving gaze shifts to generate predicted V1 firing rates. (B) Example cells from early, late, biphasic, and negative clusters, showing actual spike rasters, predicted firing rates for individual events, and normalized actual and predicted mean responses. (**C**) Actual and predicted population responses sorted by actual peak latency. (**D**) Cluster-averaged actual and predicted responses. (**E**) Top: actual versus predicted peak latency across neurons, colored by cluster type. Bottom: distribution of latency prediction error (predicted minus actual). (**F**) Cumulative distributions of median per-event residual prediction performance for true predictions and event-shuffled controls. Dashed lines indicate medians.

Trained only on head-fixed sparse noise responses, the model nonetheless predicted the relative timing of gaze shift responses, reproducing the temporal progression across the population. Predicted responses in representative early, late, biphasic, and negative cells reproduced several features of the measured gaze shift responses (Figure 5B). This correspondence was also organized across the population: when neurons were sorted by the latency of their actual gaze shift responses, the predicted responses reproduced a similar temporal progression across cells (Figure 5C; Pearson r = 0.508). Cluster-averaged responses showed the same pattern. The model captured the early positive responses most robustly, while later, biphasic, and negative responses were present but less accurately matched in amplitude (Figure 5D).

To quantify this temporal correspondence more directly, we compared actual and predicted peak latencies across cells. Predicted and measured peak latencies were strongly correlated (Spearman ρ = 0.616, p = 1.28 × 10⁻²⁹; Figure 5E), with identical median latencies of 128 ms and a median absolute error of 16 ms. Visual filters estimated under head-fixed sparse-noise conditions therefore captured the relative ordering of when neurons responded during freely moving gaze shifts.

Because each gaze shift samples a different part of the visual scene, we next asked whether the model captured not only the average response sequence, but also responses to individual events. For each neuron, we first subtracted the mean traces from both measured and predicted event traces to capture the event-by-event variation, and then measured their correlation across individual gaze shift events and compared it with an event-shuffled control, in which predicted traces were randomly paired with measured traces from different events from the same neuron (Figure 5F). Event-by-event correlations were consistently higher than the shuffled control. The median per event correlation was 0.054 for the true prediction and -0.001 for the shuffled control, and 90.7% of neurons (243 of 268 cells) showed higher correlations for the true prediction than for the shuffled control (two-sided Wilcoxon signed-rank test, p = 4.03 × 10⁻⁴¹; 2 cells were excluded because valid predicted event traces were unavailable). Although modest in magnitude, this event level correspondence links V1 responses to the specific visual input brought into view by each gaze shift.

The model recovered the response shape, relative timing, and a small but reliable component of event to event variability. It did not, however, reproduce the overall firing rate level or the negative components of some gaze shift responses during free movement: predicted peak responses were smaller than measured responses, with a median predicted to measured peak amplitude ratio of approximately 0.19. This difference in response scale may reflect behavioral state, motor-related signals, recurrent dynamics, and other factors that can elevate firing rates during free movement but were not included in the model. Together with the receptive field structure detected during gaze shifts, these results indicate that visual tuning measured under controlled conditions is informative about both the spatial organization and temporal ordering of gaze shift responses, while the overall response level depends on additional factors.

### V1 population activity shifts toward newly sampled visual input

At the single neuron level, gaze shift responses were linked to visual content through response magnitude, spatial receptive field structure, and response timing. We next asked whether this relationship was also present at the population level. We examined how decoded visual representations changed around movement onset. To do this, we trained ridge regression decoders to reconstruct shift-corrected world camera images from V1 activity during freely moving behavior. Decoders were trained on 50% of each freely moving session and evaluated on the remaining data. To relate decoding dynamics to the temporal response sequence described above, we trained decoders using either the full population or cells from each response cluster separately, within each animal. For cluster-specific analyses, decoders were trained only when at least three cells from the corresponding response cluster were sampled in a given session; sessions that did not meet this criterion were not used for decoder training.

Representative reconstructions captured coarse spatial structure in the visual scene around gaze shift onsets (Figure 6A). Decoding accuracy was concentrated in spatially localized regions rather than distributed uniformly across the image (Figure 6B), indicating that V1 population activity carried information about specific regions of the visual inputs, consistent with spatially localized receptive fields. Subsequent analyses focused on spatially localized decoding regions defined using training data within each animal.

**Figure 6.**
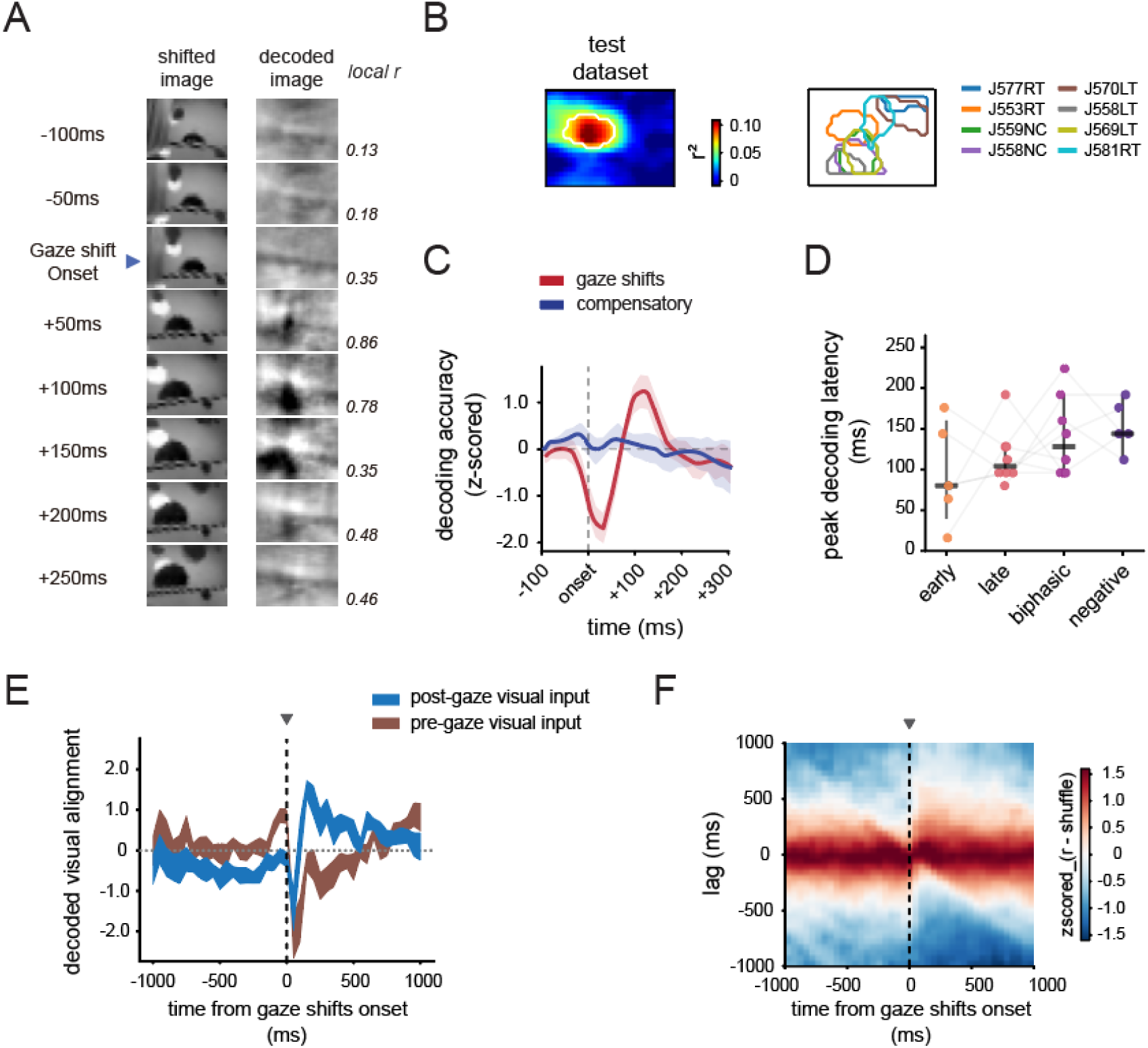
Gaze shifts temporally organize V1 population coding around newly sampled visual input. (**A**) Representative shifted images and decoded images aligned to gaze shift onset, with local Pearson correlations indicating image similarity. (**B**) Example decoding accuracy map from test dataset. White line indicates the ROI boundary for further analyses. Right, boundaries of the decoding regions identified across animals. (**C**) Mean normalized decoding accuracy for gaze shifts and compensatory movements, computed from the most reliably decoded image regions within each animal. Shading indicates SEM. (**D**) Peak decoding latency for early, late, biphasic, and negative cluster-specific decoders. Points indicate individual sessions, gray lines connect response classes within the same session, and horizontal bars indicate median latency. (**E**) Session normalized correlations between decoded images and world camera frames sampled before versus after gaze shift onset, showing a shift toward post-movement frames following each gaze shift. (**F**) Population-averaged lag-by-time map of decoded to world image similarity after shuffled baseline subtraction. Positive lags indicate frames sampled after gaze shift onset, and negative lags indicate frames sampled before it.

Using the full population decoder, decoding accuracy showed a pronounced time-locked increase after gaze shift onset, peaking approximately 100-150 ms later, whereas compensatory movements showed a relatively flat trend (Figure 6C; movement-by-time interaction: F(24,168) = 4.38, p = 5.09 × 10⁻⁹). However, mean decoding accuracy across 0 - 250 ms after the movement onset did not differ between movement types (Wilcoxon signed-rank test, p = 0.844), indicating that the V1 population carried visual information within a specific temporal window rather than uniformly throughout the post-movement period.

To ask whether this temporal organization followed the single-cell response sequence, we evaluated cluster-specific decoders over the same full-population decoding region. Median peak latency across sessions progressed from 80 ms for the early decoder to 104 ms for the late decoder, 128 ms for the biphasic decoder, and 144 ms for the negative decoder (Figure 6D). Peak latency increased across the temporal response sequence (within-session permutation test, p = 0.045). Thus, the timing of decodable visual information broadly followed the response sequence observed across single neurons.

We next asked which visual input, relative to the time of the gaze shift, this decoded activity was aligned with. For each decoded image, we measured its similarity to world camera frames sampled at different time lags relative to the same neural activity window. Following gaze shift onset, decoded images became more similar to frames sampled after the gaze shifts, while their similarity to pre-movement frames decreased (Figure 6E). The same transition was also seen at the population level (Figure 6F). To quantify this transition across sessions, we computed a lag-alignment interaction index reflecting the post-gaze increase in alignment toward later versus earlier visual frames. The index was larger for gaze shifts than for compensatory movements in all eight sessions (median paired difference = 0.0107; two-sided Wilcoxon signed-rank test, p = 0.0078; Figure S6). Applying the same analysis to the world camera movie showed that gaze shifts produced a corresponding transition towards the new visual input (Figure S7). Taken together, these results suggest that decoded V1 activity aligns to newly sampled visual input following gaze shifts.

## DISCUSSION

Across many species, vision unfolds as a sequence of active sampling events. The eyes and head are continually redirected to bring new portions of the world into view, alternating with brief stabilizing periods that hold the image relatively steady (Land 2019; Yarbus 1967). This rhythm of redirecting and stabilizing gaze has long been recognized as a core feature of active visual processing (Ahissar and Arieli 2001; Schroeder et al. 2010; Boi et al. 2017; Rucci and Victor 2015). Yet it has remained unclear how the visual changes associated with individual gaze shifts contribute to the response sequence in V1 neurons during freely moving behavior. Our results show that this sequence is not a fixed movement-aligned response. Its expression varies with the visual change delivered by each gaze shift, remains linked to each neuron’s visual response properties, and follows a consistent temporal order.

### Visual input shapes V1 neuron responses following gaze shifts

A central question in active sensing is how sensory areas represent what is sampled within the motor and behavioral context of sampling (Schroeder et al. 2010; Leszczynski and Schroeder 2019). This question is amplified by evidence that spontaneous movements account for a substantial portion of activity throughout the mouse brain, including primary visual cortex (Musall et al. 2019; Stringer et al. 2019). Our event-by-event analyses address this directly. Visual change explained more unique response variance than the measured motor variables, and such modulation was not prominently observed in darkness, when gaze movements were preserved but visual input was removed.

This is consistent with our previous finding that the gaze shift response sequence is largely eliminated in darkness (Parker et al. 2023), and extends it from the presence of the sequence to its expression on individual events. It also parallels recent work in primates showing that movement related modulation of V1 activity is largely accounted for by retinal input rather than movement alone (Talluri et al. 2023) and recent marmoset work in which gaze-related V1 responses are strongly reduced in the absence of visual input (Li et al. 2026). Across species, V1 responses during gaze shifts reflect what the movement brings into view, not simply that a movement occurred.

Further, the effect of visual change is not expressed as a uniform additive change across the V1 population. Larger visual changes were associated with enhanced positive responses in early and late cells and deeper negative responses in biphasic and negative cells. The simplest account of this pattern is that the amplitude of each response, whether positive or negative, scales with the amount of change in the visual input that corresponds to each cell’s visual tuning properties. A single global drive added to the population would shift activity in the same direction across cells, whereas scaling of each neuron’s own visually evoked response produces the bidirectional pattern we observed.

The visually evoked responses also reflected each neuron’s spatial receptive field structure. Receptive fields estimated from gaze shift epochs showed a consistent positive correspondence with those measured during head fixation, whereas compensatory epochs, which produce smaller changes in the sampled image, yielded weaker and less reliable spatial structure. Parker et al. (2022) showed that V1 receptive fields can be recovered during free movement. Here we show that this organization is carried by the spikes recruited within the brief windows of active sampling following each gaze shift. While Parker et al. (2023) established the gaze shift response sequence and its dependence on visual tuning, here we further show that this sequence encodes newly sampled visual input on an event-by-event basis.

### The sampling cycle structures the temporal cascade of V1 responses

Each gaze shift compresses a large amount of scene change into a short time window, converting spatial structure in the environment into a rapid temporal signal at the retina (Kuang et al. 2012; Rucci and Victor 2015; Intoy et al. 2024). A cortex that processes such input may therefore produce responses whose temporal ordering reflects how different image features are processed after a sample is delivered. This provides a useful framework for interpreting the cascade described by Parker et al. (2023), in which neurons tuned to coarser spatial scales respond earlier while neurons tuned to finer scales are recruited later. This progression aligns with a long-standing observation in primate V1, where the temporal evolution of spatial frequency tuning carries low-frequency information first and shifts toward higher frequencies over the response (Bredfeldt and Ringach 2002; Mazer et al. 2002), and is consistent with coarse-to-fine processing observed in mouse V1 during natural scene viewing (Skyberg et al. 2022). It also relates to work in macaque V1 showing that saccades can change how neurons represent different spatial scales of the visual scene, including a shift in sensitivity toward finer spatial structure after the movement (Niemeyer et al. 2022). Because similar coarse-to-fine dynamics are also observed during head-fixed visual stimulation, this temporal progression is unlikely to arise solely during gaze shifts. Instead, gaze shifts may provide a natural temporal input over which coarse-to-fine dynamics already present in V1 can unfold.

The visual filter analysis supports this interpretation in a direct way. Filters trained under head-fixed sparse noise conditions recovered the relative timing of gaze shift responses during free movement without being fit to gaze shift activity. The model did not fully reproduce the overall firing rate level, but it captured response shape, relative timing across neurons, and a small but reliable component of event to event variability. This dissociation suggests that visual tuning sets what visual information is represented and when it appears in neural activity following a gaze shift, whereas behavioral state, motor-related signals, and/or recurrent dynamics set how strong the response is.

### Population activity shifts toward newly sampled visual input

At the population level, an active sampling model predicts that V1 activity should change as the animal replaces one visual sample with another. Our decoding analysis supported this prediction. Decoded V1 activity became more closely aligned with visual frames sampled after gaze shift onset than with the pre-movement scene. Applying the same analysis to the world camera movie showed a corresponding transition in the visual input itself, providing a reference for interpreting the neural shift. At the level of scene content recovered by this decoder, population activity in V1 tracked the visual input sampled by each gaze shift.

The timing of this population shift also reflected the sequence observed at the single cell level. Cluster-specific decoders peaked at progressively later times after gaze shift onset, broadly matching the response timing of early, late, biphasic, and negative cells. This indicates that the gaze shift cascade is not only a sequence of firing rate changes, but also a sequence in which different subpopulations carry their strongest decodable visual information at different moments after the sample is delivered. This extends the coarse-to-fine ordering of spatial tuning across the cascade (Parker et al. 2023; Skyberg et al. 2022) from a single neuron to the visual content that can be decoded from the population.

This population result also helps frame the relationship between mouse gaze shifts and primate saccades. Receptive field remapping and predictive feedback have been documented in primate visual areas, where representations of post-saccadic locations can emerge before the eyes arrive (Sommer and Wurtz 2008). Our data do not exclude predictive components, especially in signals or brain areas not captured by this linear decoding analysis. However, at the level of decodable scene content in mouse V1, the clearest decodable pattern was an alignment with visual input sampled after gaze shift onset. This may reflect differences in visual ecology and anatomy. Mouse gaze shifts often involve large field of view updates produced by coordinated head and eye movements, and mouse V1 is more comparable to peripheral than foveal primate vision (Liska et al. 2024; Niell et al. 2026). Under these conditions, decoded V1 activity may be more strongly tied to incoming visual input than to anticipatory remapping of future scene content.

### Implications for active vision across species

Whether studied in foveate primates making rapid saccades, marmosets navigating freely with head-mounted eye tracking (Singh et al. 2025; Li et al. 2026), or afoveate mice coordinating head and eye movements during natural exploration (Meyer et al. 2020; Michaiel et al. 2020; Parker et al. 2022, 2023), visual input to cortex is structured by a sequence of active sampling events. The statistics of that input differ substantially from those produced in controlled laboratory settings (Matthis et al. 2022; Muller et al. 2023). Across these species, despite large differences in eye movement strategy and cortical magnification, the response cascade following each sample reflects an underlying coarse-to-fine processing logic (Niemeyer et al. 2022; Parker et al. 2023). Our results add to this picture by quantifying, on individual sampling events in a freely behaving animal, how the specific content of each visual sample shapes single neuron responses, population dynamics, and the temporal availability of decodable scene information.

This view also clarifies what V1 may provide for downstream regions to compute. If V1 represents each newly sampled input in a temporally ordered population response, then the integration of successive samples into a stable scene may occur further along the visual hierarchy, in posterior parietal cortex, higher visual and temporal association areas, or medial temporal lobe circuits whose activity is tightly coupled to visual exploration. Consistent with this possibility, primate entorhinal neurons represent gaze position and visual space during visual exploration (Killian et al. 2012; Meister and Buffalo 2018), hippocampal theta activity is aligned with saccadic sampling (Jutras et al. 2013), and recent work in a highly visual bird showed that hippocampal place codes can be activated by gaze toward distant locations and temporally organized around head saccades (Payne and Aronov 2025). How these samples are integrated across movements to form a stable representation of the world remains an open question at the interface of visual cortical and mnemonic processing.

## Author Contributions

JS and CMN conceived the project. CMN supervised all aspects of the project. PRLP led mouse experiments. JS led data analysis. JS, DMM, and ETTA contributed to data analysis. JS and CMN contributed to writing and editing the manuscript.

## Acknowledgements

We thank the members of the Niell Lab for helpful feedback on previous drafts of the manuscript. This work was supported by the Simons Foundation SFI-AN-NC-SCN-00007276-04 (CMN) and NIH grants R01NS127305 (CMN) and R01NS121919 (CMN).

## METHODS

All data analyzed in this study were acquired as part of the previously published datasets described in Parker et al. (2022, 2023). No new animal experiments or neural recordings were performed for the present study. Full details of animal preparation, chronic probe implantation, head-mounted camera and inertial-sensor hardware, behavioral habituation, electrophysiological recording, head-fixed visual stimuli, freely moving arena recordings, spike sorting, and histological verification are provided in those studies. Below, we summarize the aspects of the dataset directly relevant to the present analyses and provide full methodological details for the analyses introduced here.

### Experimental dataset

Recordings were obtained from eight adult C57BL/6J mice, including three males and five females, chronically implanted with silicon probes targeting the left primary visual cortex. V1 targeting was confirmed using visually evoked activity during head-fixed mapping stimuli and post hoc histology, as described in Parker et al. (2022, 2023). Mice were fitted with a recoverable head-mounted recording system that allowed repeated attachment of an eye-facing camera with infrared illumination, a wide-angle world-facing camera, and an inertial measurement unit. All procedures were performed in accordance with NIH guidelines and approved by the University of Oregon Institutional Animal Care and Use Committee.

### Recording sessions

Each recording day included head-fixed visual stimulation sessions followed by a freely moving session. During these sessions, extracellular spiking activity was recorded simultaneously with eye position, head rotation, mouse position in the arena, and head-mounted world-camera video. Head-fixed and freely moving recordings from the same day were concatenated for spike sorting so that single units could be tracked across behavioral conditions. The present analyses used only units that passed the inclusion criteria described below.

### Head-fixed visual stimulus and freely moving behavior

Head-fixed visual stimuli were presented on a monitor positioned approximately 27.5 cm from the right eye using Psychtoolbox-3. Four stimulus classes were presented during the head-fixed session: white noise, drifting gratings, contrast-reversing checkerboards, and flashed sparse noise. In the present study, responses to white noise were used to estimate head-fixed visual receptive fields for comparison with freely moving receptive fields, and responses to flashed sparse noise were used to train feature-based visual filters for predicting gaze shift responses during free movement. The flashed sparse noise stimulus consisted of circular spots of 2-32° diameter presented for 250 ms per frame with no inter-stimulus interval. Drifting gratings and contrast-reversing checkerboards were used for visual characterization and V1 verification as described in Parker et al. (2022, 2023).

During freely moving sessions, mice explored a 48 × 37 × 30 cm arena. Three walls were covered with acrylic sheets printed with high- and low-spatial-frequency gratings and white-noise patterns, and the fourth wall contained a 61 cm monitor displaying moving sparse noise. The floor was a gray silicone mat overlaid with black and white Lego bricks to provide three-dimensional visual contrast. Small tortilla chip pieces were lightly scattered to encourage foraging; animals were not food or water restricted.

### Data preprocessing

Raw recordings from head-fixed and freely moving sessions were concatenated for spike sorting, allowing single units to be tracked across the entire experiment. The global noise was removed by subtracting the median across all channels at each timepoint. Spike sorting was performed using Kilosort 2.5 (https://github.com/MouseLand/Kilosort), and single units were curated in Phy 2.0 (beta 5) based on contamination rate (<10%), mean firing rate (>0.5 Hz), autocorrelogram, and waveform shape. After sorting, data were split back into individual sessions for further analysis.

Pupil position was extracted from eye camera data (Michaiel et al. 2020). Videos were deinterlaced to 60 fps, eight points around the pupil were tracked using DeepLabCut (Mathis et al. 2018), and an ellipse was fit to compute angular pupil position. World camera data were similarly deinterlaced and corrected for lens distortion using OpenCV. Mouse position in the arena was tracked from a top-down camera via DeepLabCut. Running was defined as neck-point velocity > 2 cm s⁻¹, and stationary periods as < 2 cm s⁻¹.

Horizontal head rotation velocity was extracted from the IMU, converted to deg s⁻¹, and interpolated to eye camera timestamps. Leftward and rightward directions were defined relative to the animal’s perspective. Head movements exceeding 60 deg s⁻¹ were classified as gaze shifts or compensatory movements based on gaze velocity, defined as the sum of horizontal eye and head velocities (Michaiel et al. 2020). High gaze velocity (> 240 deg s⁻¹) indicated gaze shifts, low velocity (< 120 deg s⁻¹) indicated compensatory movements, and intermediate values were excluded. Only movement onsets were used for event-aligned analyses, and compensatory movements occurring within 500 ms of any gaze shift were excluded from analyses comparing gaze shift and compensatory epochs.

To account for continuous changes in eye and head position during freely moving behavior, world camera frames were corrected for gaze-induced image shifts using the Shifter Network (Parker et al. 2022). This network takes instantaneous eye and head position as input and applies a spatially uniform shift to each frame to approximate the retinal image at a common reference gaze angle, enabling comparison of visual input across time points with different gaze positions. All subsequent analyses involving world camera images used these shift-corrected frames.

### Cell inclusion criteria for the present analyses

The main analyses focused on 270 gaze shift responsive V1 units from the eight paired head-fixed and freely moving recording sessions. Cells were included if they had a mean firing rate of at least 1 Hz in the head-fixed visual stimulus session, a mean firing rate of at least 1 Hz during free movement, a mean gaze-shift-aligned firing rate of at least 1 Hz, and an absolute gaze response modulation index of at least 0.1. Cells were also required to have a non-noise gaze shift response cluster label. The response groups were early (n = 51), late (n = 77), biphasic (n = 90), and negative (n = 52) cells. Unless otherwise stated, visual transient modulation, receptive field estimation, and head-fixed visual filter prediction analyses used this same 270 cell pool.

For analyses with additional technical requirements, the same cell pool was further restricted only when necessary. Population decoding analyses were performed separately within each animal using the subset of simultaneously recorded units assigned to the gaze shift responsive clusters.

### Visual transient modulation of V1 gaze shift responses

Visual change associated with each gaze shift was quantified from the shift-corrected world camera images. For each event, we measured luminance and spatial-frequency transients during the 0-250 ms period after gaze shift onset. Luminance change was quantified from the magnitude of positive and negative frame-to-frame intensity changes across the image. To quantify spatial frequency change, frame-to-frame difference images were transformed using a two-dimensional Fourier transform, and spectral power was measured within low (0-0.10 cycles/pixel), middle (0.10-0.20 cycles/pixel), and high (0.20-0.35 cycles/pixel) spatial frequency bands. The peak luminance and spatial frequency changes during the post-gaze window were standardized across events within each session and averaged to obtain a single visual transient index per event. Events were divided at the within-session median for high- and low-visual-change comparisons, whereas the continuous index was used for regression analyses.

Neural responses were baseline-subtracted using the −500 to −200 ms period before gaze shift onset. For each cell, the mean response during the 0-250 ms post-gaze period was separated into positive and negative responses. Individual event responses were projected onto these components to obtain event-wise positive and negative response amplitudes, allowing mixed-sign response profiles to be quantified without cancellation.

To estimate the contributions of visual and motor variables to event-by-event response variability, positive and negative response amplitudes were modeled using the visual transient index, head amplitude, eye amplitude, and running speed. Predictors were standardized within each session. Model performance was evaluated using five-fold cross-validation for cells with at least 60 valid events. The unique contribution of each predictor was calculated as the difference in R² between the full model and a reduced model in which that predictor was omitted, using the same cross-validation folds.

For event-by-event visual transient analyses, baseline-subtracted firing rate at each peri-gaze time bin was regressed across events against the standardized visual transient index, yielding a visual modulation slope for each cell. Positive and negative modulation were summarized within the 0-250 ms period after gaze shift onset.

As an independent motor control, we analyzed paired light and dark freely moving recordings collected from the same animal and recording day with matched unit sorting. Cluster identity and preferred gaze direction were defined from the light session and applied to the corresponding dark session, and gaze shifts were detected using the same movement criteria. Dark gaze shifts were divided into low- and high-amplitude groups using a within-session median split of total gaze amplitude. In parallel, baseline-subtracted firing rate was regressed at each time point against z-scored gaze amplitude from −200 to 300 ms.

### Estimating visual receptive fields

Visual receptive fields were estimated for head-fixed and freely moving conditions using a regularized linear model with temporal lags and L2 regularization, consistent with prior receptive field mapping approaches (Pillow et al. 2008). Binned spike responses were modeled as a linear function of temporally shifted stimulus frames, and receptive field weights were estimated by solving a regularized least squares problem:

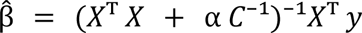

where β^ denotes the receptive field weights, C⁻¹ is the regularization matrix, and α controls regularization strength. The regularization matrix included a spatial smoothness prior over neighboring pixels, with the intercept term left unregularized. The regularization strength was selected from a logarithmically spaced grid of α values by minimizing mean squared error on held-out test data at zero temporal lag. The selected α value was then reused across temporal lags for the same cell. For gaze shift and compensatory receptive field estimates, the α value selected from active freely moving data was reused.

Stimulus frames were corrected for eye and head movements using the Shifter Network (Parker et al. 2022), sampled at 16 ms resolution, and cropped to 20 by 30 pixels to remove edge artifacts. Receptive fields were estimated across a grid of temporal lags, and each receptive field map was reconstructed by reshaping the fitted spatial weights to the dimensions of the cropped stimulus. Temporal shifts were applied without circular wrapping, so samples that would fall outside the recording boundaries were excluded.

For freely moving analyses, receptive fields were estimated from active periods of freely moving behavior, gaze shift epochs, and compensatory movement epochs. Gaze shift and compensatory epochs were defined as peri-event windows spanning −50 to 250 ms. To reduce contamination from nearby gaze shift events, compensatory events occurring within 500 ms of any gaze shift were excluded before fitting. Because the number and temporal overlap of peri-event frames can differ between movement types, gaze shift and compensatory analyses were matched by the number of frames used for receptive field fitting.

A cell was considered to have significant receptive field structure if the training and test receptive fields were more similar to each other than expected from 1,000 control comparisons in which one map was temporally shifted relative to the other (10s to 60s). To quantify receptive field similarity across conditions, Pearson correlation coefficients were computed between receptive fields estimated during head-fixed white noise stimulation and those estimated during freely moving behavior. These correlations were used to compare the spatial correspondence of gaze shift and compensatory receptive fields with independently measured head-fixed receptive fields.

### Sparse noise trained spike prediction during freely moving gaze shifts

Visual filters were trained on responses recorded during head-fixed sparse flashing noise stimulation and applied without modification to freely moving visual input to predict gaze shift aligned spiking activity. World camera frames were cropped to 20 × 30 pixels and standardized across time. Spatiotemporal visual features were computed by dividing each frame into 5 × 10 non overlapping spatial bins (4 × 3 pixels each), yielding spatially resolved ON, OFF, ΔON, and ΔOFF features per bin. ON and OFF components were defined as the positive and negative portions of the luminance signal, respectively, and their temporal derivatives (ΔON, ΔOFF) were computed from causal frame-to-frame differences after mild spatial smoothing with a Gaussian kernel (σ = 0.75 pixels). Each spatial feature was projected onto 6 raised-cosine temporal basis functions spanning a 0 to 250 ms causal history window. To reduce parameter dimensionality and support cross-condition generalization, feature weights were parameterized as separable spatial-by-temporal filters. Poisson GLMs were fit to spike counts from the head-fixed sparse noise session using L-BFGS optimization with L2 regularization. Regularization strength was selected independently for each cell by grid search over 13 logarithmically spaced values (0.003 to 3000), evaluated on a validation set consisting of 20% of the sparse noise recording, split into contiguous 10-second chunks to avoid temporal leakage between training and validation sets.

The fitted models were applied directly to shift-corrected world camera frames from the freely moving session to generate predicted firing rates across the entire recording. Predicted and observed firing rates were then aligned to gaze shift onset. Prediction performance was evaluated on a test set of 25% of gaze shift events that was excluded from all model fitting and hyperparameter selection. Performance metrics included shape correlation between baseline subtracted predicted and actual firing rate traces, peak latency, amplitude ratio, and R² within the 16 to 250 ms post-gaze window.

For event-by-event prediction analysis, actual and predicted firing-rate traces were extracted for each individual gaze shift event. To remove correlation arising from the common gaze-aligned response profile, the mean event-aligned trace for each cell was subtracted from the corresponding actual and predicted event traces. Pearson correlation was then computed between the resulting actual and predicted residual traces within the 0-250 ms post-gaze window, and the median per-event correlation was used as the prediction score for each cell. For the shuffled control, predicted residual traces were randomly reassigned across gaze shift events within the same cell before computing the same metric, thereby disrupting event identity while preserving the residual response distributions. This procedure was repeated 1,000 times per cell to generate the shuffled distribution.

### Visual decoding using ridge regression

To evaluate how neural population activity represented visual input, we trained ridge regression decoders to reconstruct shift-corrected world camera images from simultaneously recorded V1 spike counts, following population decoding frameworks established in visual cortex (Naselaris et al. 2009; Stanley et al. 1999). For each animal, the full decoder used all neurons assigned to the four gaze shift response clusters, and cluster-specific decoders were trained separately using early, late, biphasic, or negative cells. The relationship between neural activity and visual input was modeled as a linear system with L2 regularization:

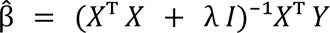

where *X* is the spike-count matrix, *Y* is the world camera image matrix, β^ is the decoder weight matrix, *I* is the identity matrix, and λ is the regularization parameter. Neural predictors were standardized using training data only, and λ was selected from a logarithmically spaced range using five-fold cross-validation within the training data.

Training and test data were separated using a temporal block-based procedure to reduce the influence of local temporal correlations. Each session was divided into consecutive 10-s blocks, with 50% of the blocks randomly assigned to the training set and the remaining blocks reserved for testing. Decoder fitting and parameter selection used only active freely moving frames within the training blocks. Held-out gaze shift and compensatory events were included only when the entire peri-event window fell within test blocks, ensuring that event-aligned decoding was evaluated on data not used for model fitting.

Spatial decoding regions were defined independently of the test data. Training blocks were divided into two subsets, and pixel-wise decoding maps were cross-fitted between the two subsets. A connected region centered on the peak of the cross-fitted training map was selected as an ROI and then used for subsequent analyses. The same training-defined region was used for gaze shifts, compensatory movements, and cluster-specific decoders. Spatial decoding performance on active test frames was quantified using pixel-wise R² between predicted and actual world camera images.

### Gaze aligned decoding analysis with temporal lag

To examine the temporal relationship between decoded V1 activity and visual input, decoded images at each gaze-aligned neural time point were compared with shift-corrected world camera frames sampled at different temporal lags. Similarity was quantified as the spatial Pearson correlation between the decoded and world camera images. Positive lags corresponded to visual frames sampled after the neural activity window, whereas negative lags corresponded to frames sampled before it. Correlations were averaged across events to generate a lag-by-time matrix for each session.

To remove correlation structure unrelated to event-specific visual content, the same analysis was repeated after shuffling event identities, and the shuffled correlation was subtracted from the true correlation. Lag asymmetry at each gaze-aligned time point was defined as the mean shuffle-corrected correlation across positive lags minus that across negative lags. A lag-alignment interaction index was then calculated as the mean asymmetry during the 0-300 ms post-gaze period minus that during the −300-0 ms pre-gaze period. Session-level indices were compared using two-sided paired Wilcoxon signed-rank and sign-flip tests. This analysis was applied to both gaze shifts and compensatory movements.

As a control for temporal structure in the visual input itself, the same lag analysis was performed directly between world camera frames. For each event-aligned time point, the current world camera frame was correlated with frames sampled at the corresponding positive and negative temporal lags, and the same lag-alignment index was calculated.

## Statistical analysis

Statistical analyses were performed using two-sided nonparametric tests unless otherwise specified. Paired and unpaired comparisons used Wilcoxon signed-rank and rank-sum tests, respectively, and comparisons across multiple response clusters used Kruskal-Wallis tests followed by post hoc pairwise tests. Population-level analyses were performed on session-level summary values rather than treating individual cells or events as independent samples. Two-sided sign-flip tests were used as an additional paired session-level test where indicated. Multiple comparisons were controlled using false discovery rate correction when appropriate. Decoding time courses were analyzed using two-way repeated-measures ANOVA with movement condition and peri-event time as within-session factors. Data are reported as mean ± SEM unless otherwise specified, with statistical significance defined as α = 0.05.

## Supplemental Figures

**Figure S1.**
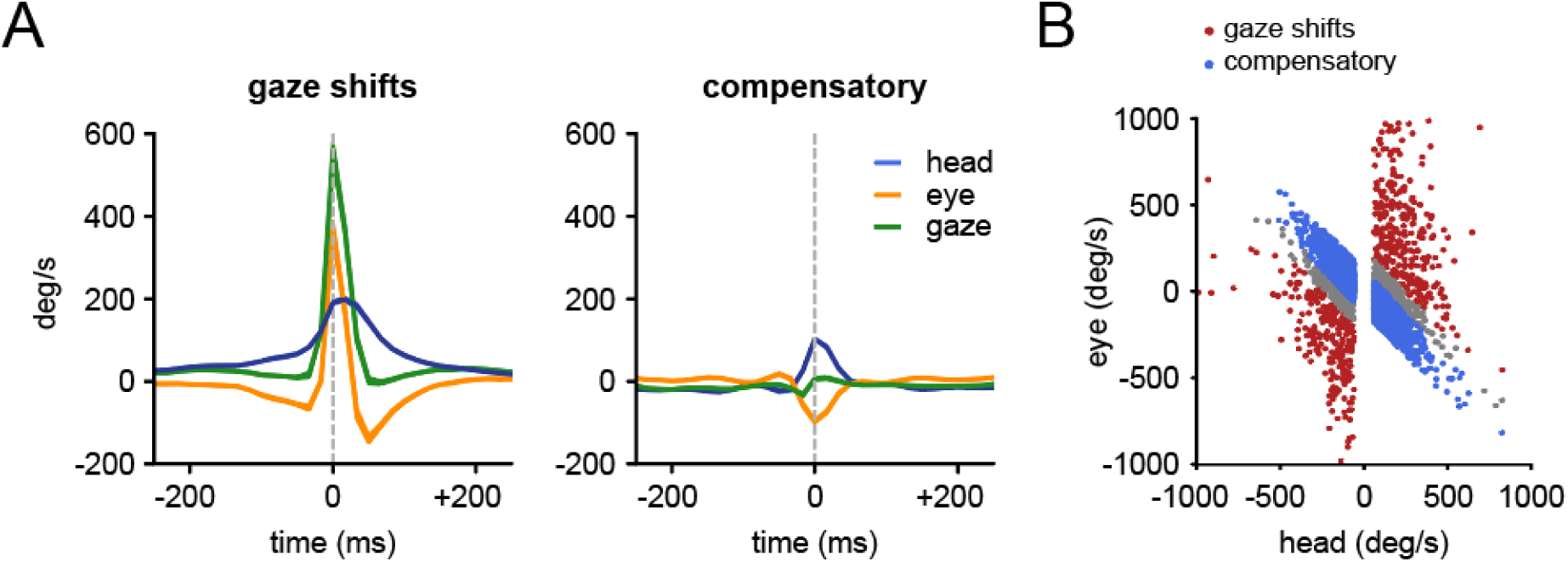
Gaze shift and compensatory eye-head movements (related to Figure 1). (**A**) Averaged head, eye, and gaze velocity traces aligned to gaze shifts and compensatory movements. Gaze shifts were associated with larger, more coordinated transients in head and eye velocity, whereas compensatory movements showed opposing eye-head dynamics and smaller net gaze change. (**B**) Distribution of eye versus head velocity samples for gaze shifts and compensatory movements.

**Figure S2.**
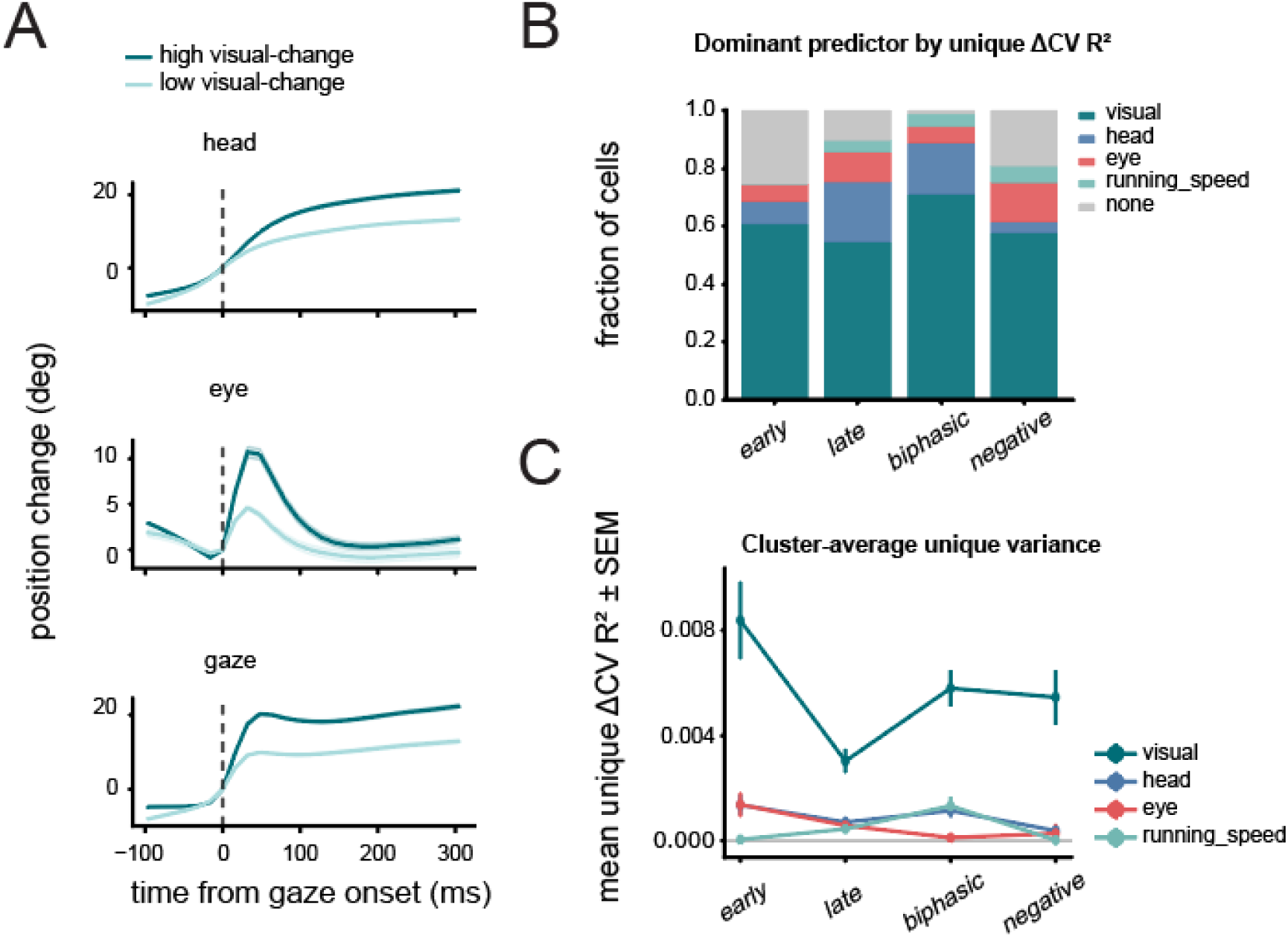
Visual transients predict V1 response modulation beyond motor covariates (Related to Figures 2 and 3). (**A**) Mean head, eye, and gaze position changes for gaze shifts with low or high visual change, defined by a session-wise median split of the post-gaze visual transient index. (**B**) Dominant predictor for each cell, defined as the predictor with the largest unique ΔCV R². Visual transient index was the most frequent dominant predictor in early (60.8%), late (54.5%), biphasic (71.1%), and negative (57.7%) clusters. (**C**) Cluster-averaged unique predictive contribution of visual transient index, head amplitude, eye amplitude, and running speed. Values show mean ΔCV R² ± SEM across cells.

**Figure S3.**
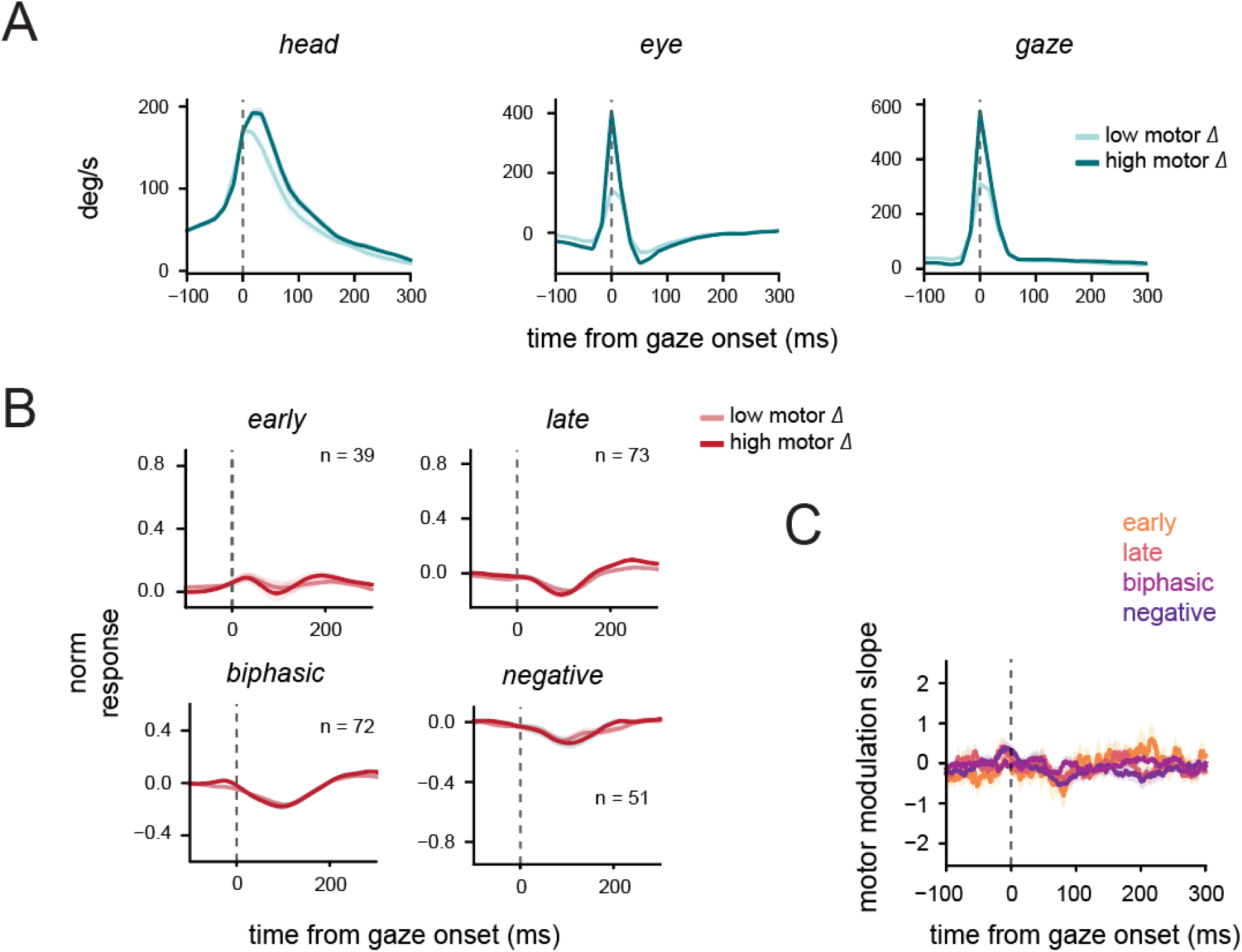
Gaze shift responses in V1 during dark sessions (related to Figures 2 and 3). (**A**) Direction-aligned head, eye, and gaze velocity traces for low-and high-amplitude gaze shifts during dark sessions. Events were divided by a median split of total gaze amplitude. Shading indicates SEM across sessions. (**B**) Normalized V1 responses in darkness for low- and high-gaze-amplitude events. Cluster identity and each cell’s preferred gaze direction were assigned from the paired light session and applied to the corresponding dark session. Shading indicates SEM across cells, and n values indicate the number of cells in each cluster. (**C**) Gaze-amplitude modulation of V1 responses in darkness. For each cell, baseline-subtracted firing rate at each time bin was regressed against the z-scored gaze-amplitude index across events, and slopes were averaged across sessions within each cluster.

**Figure S4.**
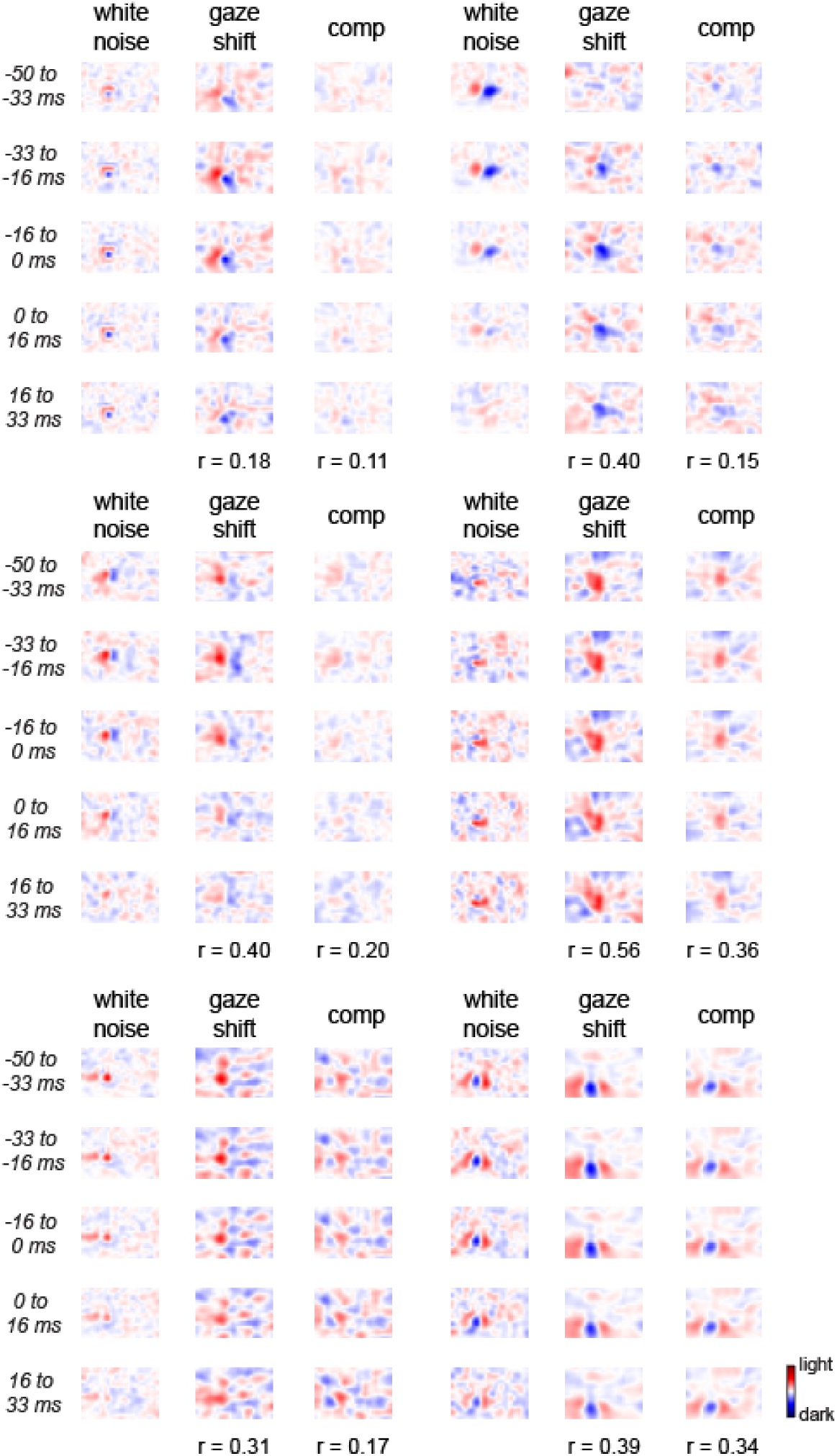
Comparison of receptive field estimates across different conditions (related to Figure 4). Spatiotemporal receptive fields estimated from head-fixed white-noise stimuli, gaze shifts, and compensatory movements for individual units.

**Figure S5.**
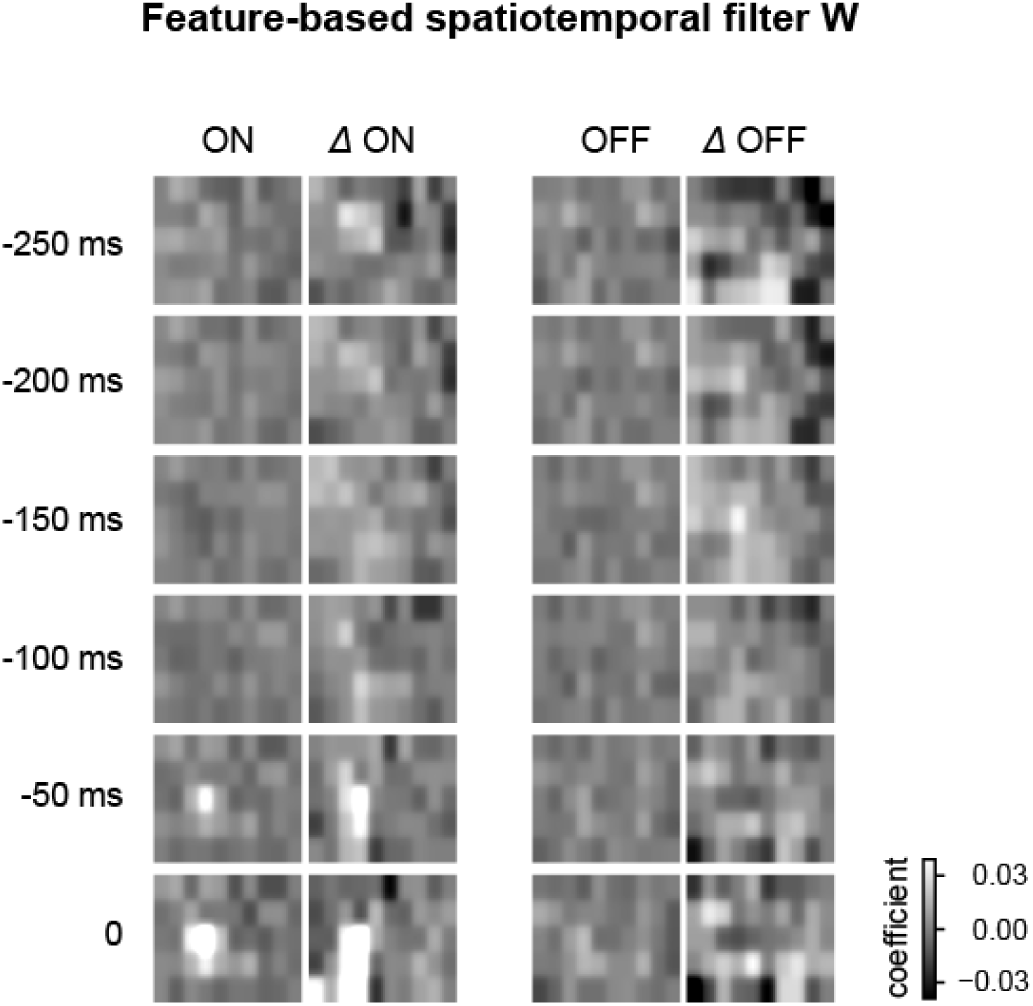
An example of the estimated visual filter (related to Figure 5). Example feature-based spatiotemporal filter estimated from head-fixed sparse-noise responses. Columns show ON, ΔON, OFF, and ΔOFF feature weights, and rows show temporal lags relative to spiking.

**Figure S6.**
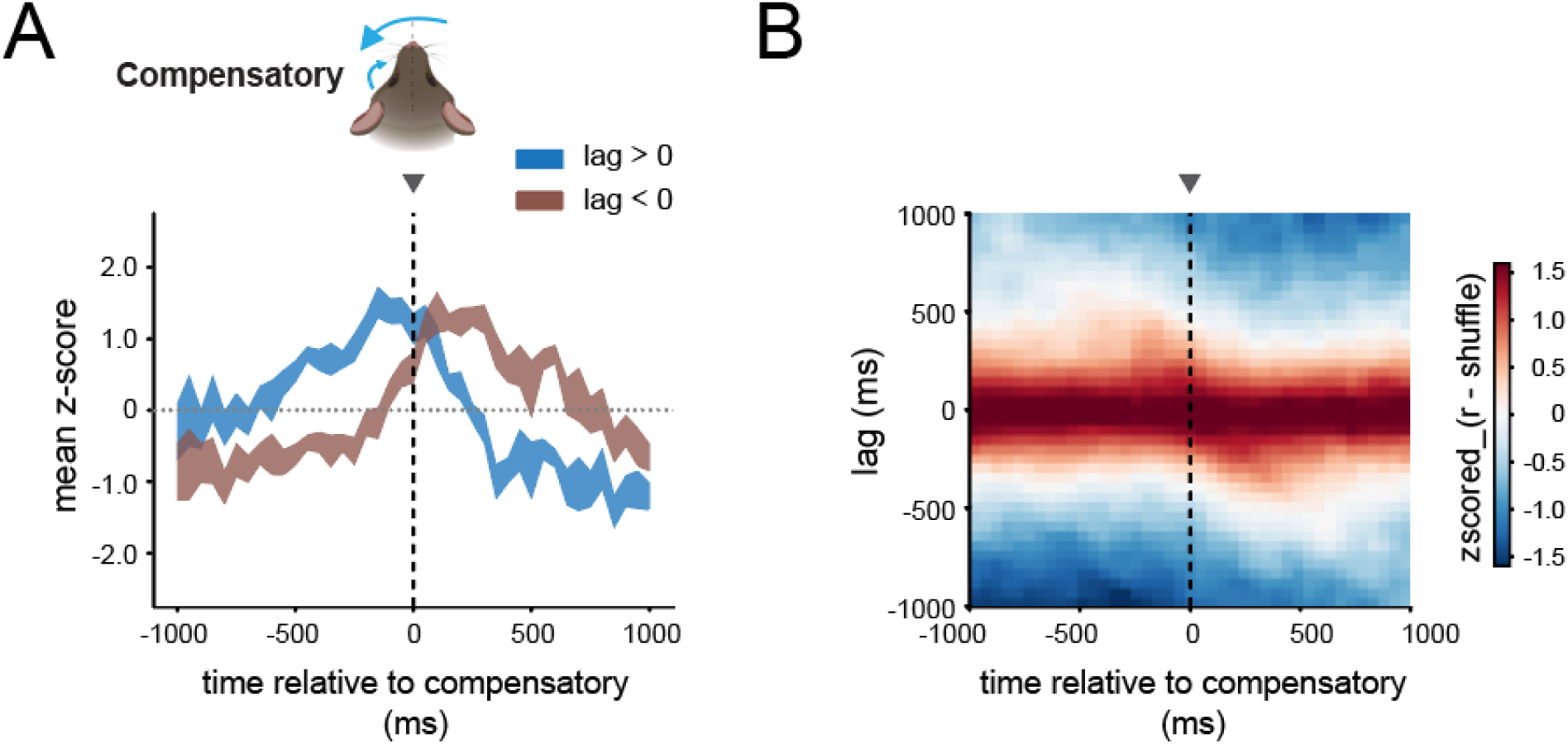
Temporal alignment between decoded V1 representations and visual input during compensatory movements (related to Figure 6). (**A**) Mean correlations between decoded V1 representations and visual frames sampled at positive and negative frame lags around compensatory movements. Unlike gaze shifts, compensatory movements did not show a clear shift toward visual input sampled after movement onset. (**B**) The corresponding population-averaged correlation map. Correlations remained largely symmetric across positive and negative visual-frame lags after subtraction of the event-shuffled baseline and normalization within each session, consistent with a more stable visual reference during compensatory movements.

**Figure S7.**
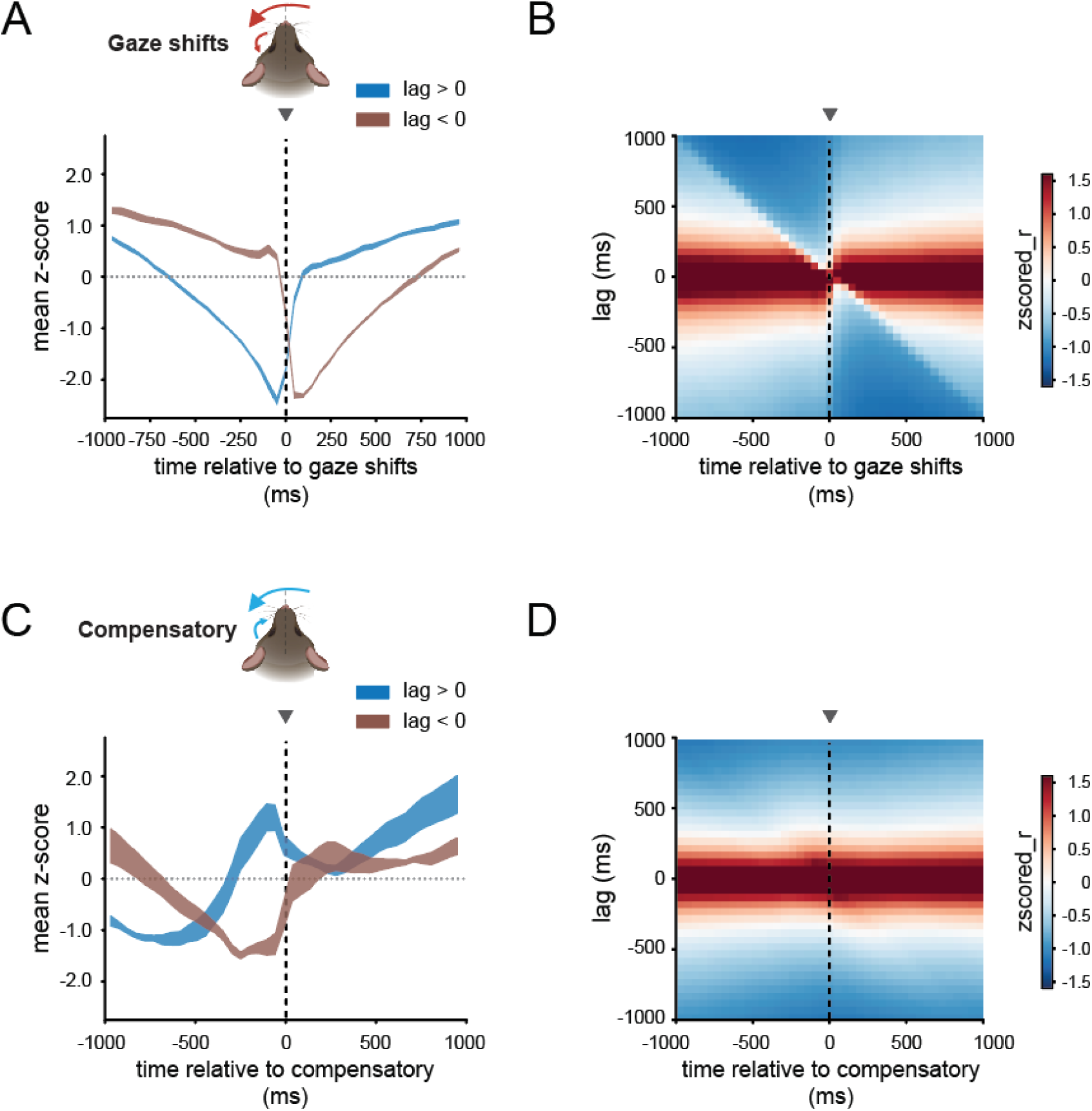
Temporal structure of world camera input around gaze shifts and compensatory movements (related to Figure 6). (**A**) Mean z-scored correlations between world-camera frames sampled at positive and negative frame lags around gaze shifts. Frame correlations showed a pronounced disruption around gaze shift onset. (**B**) The corresponding time-by-frame-lag correlation map showed a tilted structure and transient disruption of visual continuity around gaze shift onset. (**C**) Same analysis for compensatory movements. Frame correlations changed more smoothly around movement onset, without the abrupt disruption observed during gaze shifts. (**D**) The corresponding time-by-frame-lag correlation map showed a more horizontal and symmetric structure, consistent with greater visual stability during compensatory movements.

